# Roadmap to Help Develop Personalized Targeted Treatments for Autism as a Disorder of the Nervous Systems

**DOI:** 10.1101/2020.06.01.127597

**Authors:** Elizabeth B Torres

**Author notes:** Correspondence.: (011) (E.B.T).

## Abstract

There is a disconnect between the clinical behavioral definition of autism and the genomic science that this definition largely informs and steers. But the digital sensor revolution paired with open access to genomics data has the potential to bridge the gap between these two layers of knowledge. Here we use the SFARI genes module and interrogate the human genome upon removing those genes. We then compare the remaining genes’ expression on tissues responsible for brain, heart and organs function to its counterpart in well-known neurological disorders of genetic origins. Despite clinical criteria emphasizing a behavioral definition of Autism, over a neurological one, here we find convergence between Autism and the neurological disorders. Tissues involved in motor control, emotions and memory are the most affected by the removal of the SFARI Autism genes. Congruent with this picture, the Ataxias, Parkinson’s disease and Fragile X share 76.9% of the most affected tissues, including those related to motor control and autonomic function, while mitochondria disorder share 61.5% with autism. Together, these results offer a new roadmap to help diagnosis and personalized targeted treatments of autism. They underscore Autism as an objectively quantifiable disorder of the nervous systems.

---

Autism is an umbrella term defining a highly heterogeneous disorder that encompasses several important interacting axes. These axes range from aspects of the disorder that remain hidden to the naked eye, *e.g.* the microbiome, the metabolome and the genome, to observable aspects of behaviors that are currently used to define diagnosis and treatment criteria [1–3]. One facet of autism that could be both observed and physically quantified is the evolution of the *disorders of the nervous system* as the person ages and transitions to adulthood [4]. We can indeed leverage the wearable sensors revolution, digitize current inventories, and significantly improve diagnostics criteria by adding an unprecedented level of granularity to current subjective methods [5] (Figure 1A). This approach to autism, combined with genomics would be amenable to help stratify autism into subtypes and build personalized treatments, tailored to the most prominent features of each group, under the Precision Medicine (PM) model [6, 7]. The PM model can be adapted to autism with the aim to highlight the inherent capacity for social readiness of each child, using information directly related to fundamental levels of neuromotor control that escape the naked eye of the diagnostician or therapist. Such information would be necessary to aid scaffolding from the start of life, aspects of social reciprocity, mirroring, turn-taking, and other basic elements underlying social interactions and communication that fundamentally depend on the micro- and macro-movements and on the biorhythms they produce [8, 9].

**Figure 1.**
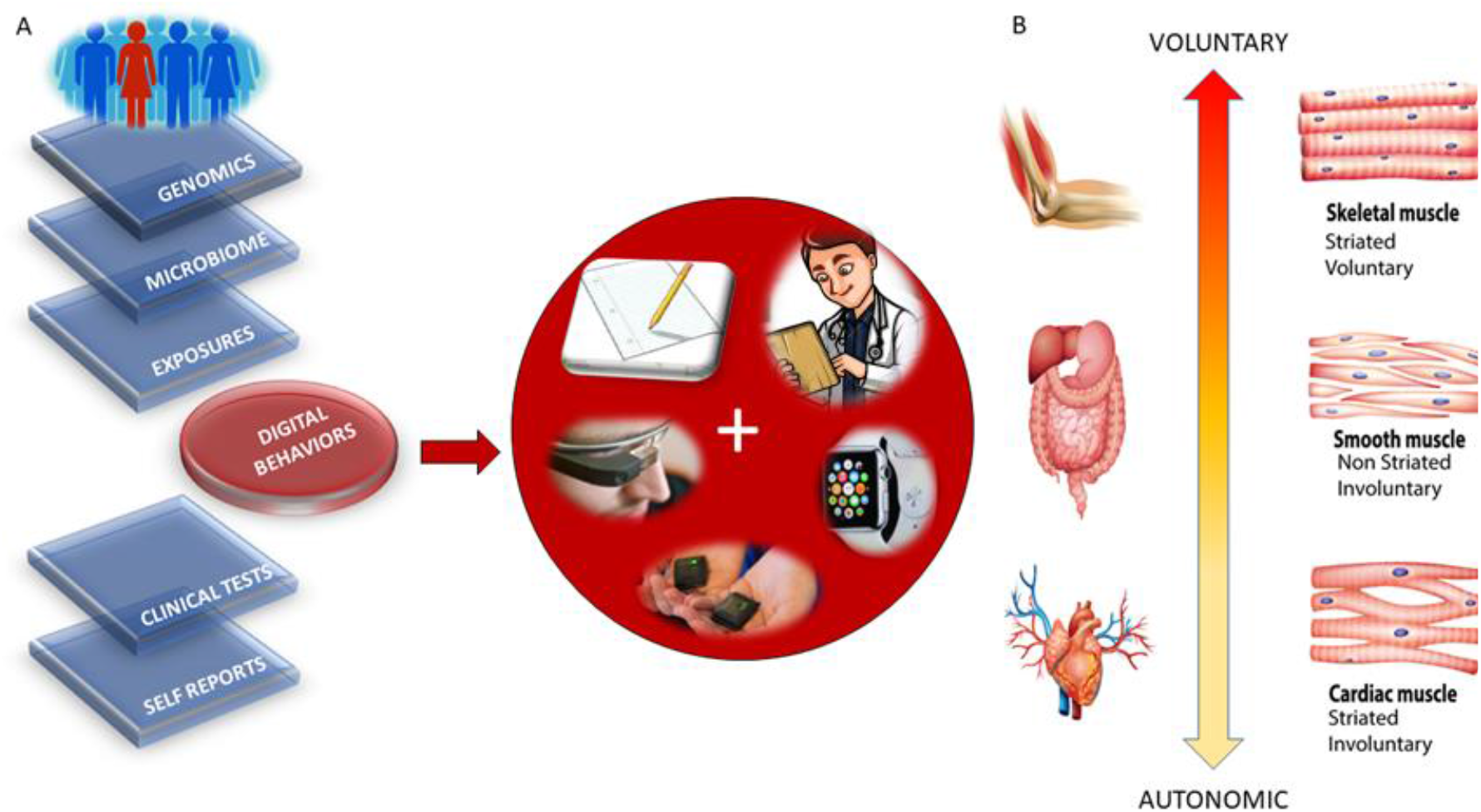
Roadmap to implement the Precision Medicine Model for ASD research, diagnoses and treatments. (A) Interconnected knowledge network can contribute information about the individual’s medical history, behaviors, environment, microbiome, and genetic makeup. Importantly, the new proposed layer of *digitized behaviors* leveraging the wearable biosensors revolution can transform medicine by creating truly personalized assessments. In Autism, this transformation could lead the way to targeted treatments. (B) A taxonomy of the nervous systems functioning based on a phylogenetic orderly maturation scale can help stratify autism subtypes and connect behaviors to genomics. By linking the fundamental muscle types to the levels of control in the nervous systems, digital behaviors can be mapped to neuromotor control levels; neuromotor control levels mapped to muscle types and muscle types to genes/proteins. Any measure of treatment’s effectiveness for ASD can then map back to improvements in behaviors embedded in activities of daily social life.

In recent years, our lab has proposed the use of PM to help reveal underlying mechanisms of various subtypes of autism, defined by a phylogenetically orderly taxonomy of the nervous systems’ functions [10], according to levels of maturation [11] (Figure 1B). This taxonomy proposes three fundamental levels of neuromotor control ranked by order of stabilization and maturation in the nervous system’s biorhythmic activities: autonomic, involuntary, and voluntary. These levels can be conceptualized as intrinsically related to three types of muscles: cardiac, smooth, and skeletal, respectively. Each one of these levels, which depend on specific proteins and genes, would then be selectively characterized, and mapped to the affected autism subtypes. The functional evolution of this affection of the nervous systems could be at least partly tracked through different tissues commonly used to characterize gene expression across the human genome. In this way, we would seamlessly connect behavioral criteria with nervous systems function and in turn, link affected tissues to their genomic underpinnings.

The advent of open access genomic databases allows us to investigate these relationships between tissues and gene expression in autism and in other disorders of the nervous systems that may or may not go on to receive an autism diagnosis; but that nonetheless present similar phenotypic features at some stage of the disorder throughout the person’s lifespan. In the past, some of these conditions would be rendered *comorbid* to the core descriptions of socially inappropriate behaviors, problems with communication, or repetitive-ritualistic movements. Yet, mounting evidence from the last couple of decades of Neuroscience research has clearly painted a more accurate picture rendering the “*autistic behaviors*” as the tip of the iceberg in a model that highlights deficits in expected social performance, but says nothing about the autistic systems’ inherent capacity for social readiness. Indeed, the current model of autism does not provide a roadmap to address the medical conditions that we see evolving in cross sectional and in longitudinal studies involving different ages [5, 12].

An age-dependent approach to autism is important. Undeniably, recent evidence suggests that aging in autism is different than typical aging in the precise sense that the nervous systems seem to be changing at an accelerated pace [12]. By 40 years of age, it is estimated that 20% of individuals manifest symptoms of Parkinsonism that seem to start at a very early age in relation to neurotypical aging systems [13]. Yet, because of the pervasive insistence that autism is a behavioral problem, these neurological conditions receive no neuroprotective therapies of any kind. In fact, the current Psychological detection criteria explicitly exclude any somatic sensory motor issues from the diagnosis [2, 14], while the Psychiatric criteria do explicitly exclude any of the motor issues [3] (and see Supplementary Material).

Autism is not currently defined as a neurological movement / sensory disorder, but in the 1970’s several neurological models competed with the then nascent behavioral ones [15–17]. Over time, the behavioral approach was well organized and gained popularity among parents and practitioners. The neurological model was abandoned in favor of behavioral modification approaches [18]. This evolution established a business model eventually sheltered by social policies that today implement and sustain the autism diagnoses and treatments [19, 20]. Over time however, it has become evident that with excess undesirable involuntary motions [12, 21, 22], a prevalence of balance (Ataxia) issues from an early age [23–26] and the presence of general issues with the autonomic [27–31] and the voluntary nervous systems [11, 32–38], it may be critical to re-evaluate the importance of seriously considering the neurobiology underlying those observed “*modifiable autistic behaviors*”. Increasing evidence pointing to somatic sensory motor issues [11, 21, 24, 25, 39–49] suggests that it may be more relevant at this point in the history of Autism, to identify when this neurodevelopmental disorder turns into a neurodegenerative one, than to continue to treat it as socially inappropriate *behaviors*.

Since many of these neurological and medical issues can be traced back to the genes and their stochastic interactions, here we take the genomic information from the general population and combine it with the genomic information derived from the autistic population-*as defined by the current clinical criteria that sidelines the neurology of Autism*. We then re-examine the general population information while considering other disorders that also present some of the autistic neurological traits and that in many cases, have high penetrance in Autism. We ask if we may arrive at convergent neurobiology pointing back to the neurological issues of the nervous systems. More specifically, we combine genetic information from the Ataxias, Fragile X related disorders, X-chromosome, Mitochondrial disorders and Parkinson’s disease to compare in 54 of the commonly tracked human tissues (including those from the brain, the heart and other organs) the outcome of removing the SFARI Autism genes from the neurotypical tissue gene expression *vs*. removing the genes known to cause the other well established neurological conditions.

## Results

The maximally affected tissues upon genes removal, according to the genes’ stochastic expression (count in Transcripts Per Million, TPM) are depicted in Table 1. This tissues’ gene expression was modelled by the exponential distribution *y* = *λe*^−*λx*^, with *x* as the genes’ combination expressed in the tissues, and *λ* as the exponential rate parameter. The Δ*λ* between the neurotypical case (containing all genes) and the disorder case (upon removal of the genes) provides a sense for the departure from the normative case. This difference, taken for the removal of the SFARI genes, is depicted in Figure 2A, with samples of maximally affected tissues in Figure 2B that are known to be critical for motor control, regulation, adaptation/learning, and coordination. The Δ*λ*-median ranking quantified the difference between neurotypical tissues’ gene expression *vs.* tissues’ gene expression upon removal of the genes corresponding to the disorders in question, with 4 groups ordered by the size of Δλ. This *λ*-quantity was first obtained relative to the neurotypical population tissues, *i.e.* including all the counts (gene expression) from all genes, *i.e.* to model an exponential process. The computation of *λ* using maximum likelihood estimation (MLE) is explained in the Appendix A. It does not assume any order of the counts, but rather seeks to identify the resulting *λ* for each tissue, treating the genes’ count (expression) as a random, memoryless stochastic process. Typically, the exponential distribution is used to model times between events, but here we used it to model the fluctuations in the values of the counts across the genes as they randomly fluctuate their expression across each of the 54 tissues. We note this to underscore that the results spontaneously self-emerge from the random combination of the genes involved, rather than from the clinical criteria used to denote the genes’ relevance to Autism. There is in fact no scoring of such relevance for the genes associated to the other neurological disorders under consideration (*e.g.* the Ataxias, Parkinson’s, etc.)

Removal of the Autism SFARI genes affected all tissues related to brain areas known for motor control, motor learning and adaptation, and motor coordination. They also affected brain regions required in memory (hippocampus) and emotion (amygdala) and tissues important for systemic organs’ functioning such as those containing smooth muscles, cardiac and skeletal muscles in the taxonomy proposed in Figure 1B. Furthermore, in 10/13 (76.9%) of the tissues, they overlapped with the tissues affected by removing the combined genes from the well-known neurological disorders. Supplementary material Table 1 shows the breakdown of the most affected tissues revealed from selectively removing from the human genome, the genes from each of these neurological disorders separately. The results for the top ranked group of tissues are congruent with the results obtained from the combined gene removal using the genes from the Ataxias, X, FX and PD (albeit shuffling tissues order when breaking them down by the SFARI scoring). The Mitochondrial disease case however shows 8/13 (61.5%) congruence of the most affected tissues and higher ranking for heart related tissues.

**Table 1.** Highest affected tissues upon removal of the SFARI genes linked to Autism from the human GTEx database (column 1). Results from the removal of the combined genes in the Ataxias, X-Chromosome, Fragile X, Parkinson’s disease and Mitochondrial disease (column 2): shaded are 10/13 tissues (76.9%) of the top median ranked tissues maximally affected in Autism and 100% concurrent with those affected in the other neurological conditions of known genetic origins. Column 3 is the same as in column 1 while removing from the SFARI Autism set 14 genes that overlap with the Ataxias and PD (see those genes listed in Supplementary Table 2).

| SFARI Autism | Ataxias, X, FX, PD, Mitochondria | SFARI Autism without overlapping Ataxia, PD genes |
| --- | --- | --- |
| Putamen Basal Ganglia | Hippocampus | Putamen Basal Ganglia |
| Substantia Nigra | Amygdala | Substantia Nigra |
| Amygdala | Substantia Nigra | Amygdala |
| Hippocampus | Heart Left Ventricle | Hippocampus |
| Caudate Basal Ganglia | Whole Blood | Caudate Basal Ganglia |
| Anterior Cingulate Cortex | Muscle Skeletal | Nucleus Accumbent Basal Ganglia |
| Nucleus Accumbent Basal Ganglia | Nucleus Accumbent Basal Ganglia | Anterior Cingulate Cortex |
| Hypothalamus | Putamen Basal Ganglia | Hypothalamus |
| Brain Cortex | Anterior Cingulate Cortex | Brain Cortex |
| Frontal Cortex | Brain Cortex | Frontal Cortex |
| Heart Left Ventricle | Hypothalamus | Spinal Cord |
| Spinal Cord | Heart Atrial Appendage | Heart Left Ventricle |
| Pancreas | Caudate Basal Ganglia | Kidney Cortex |

**Figure 2.**
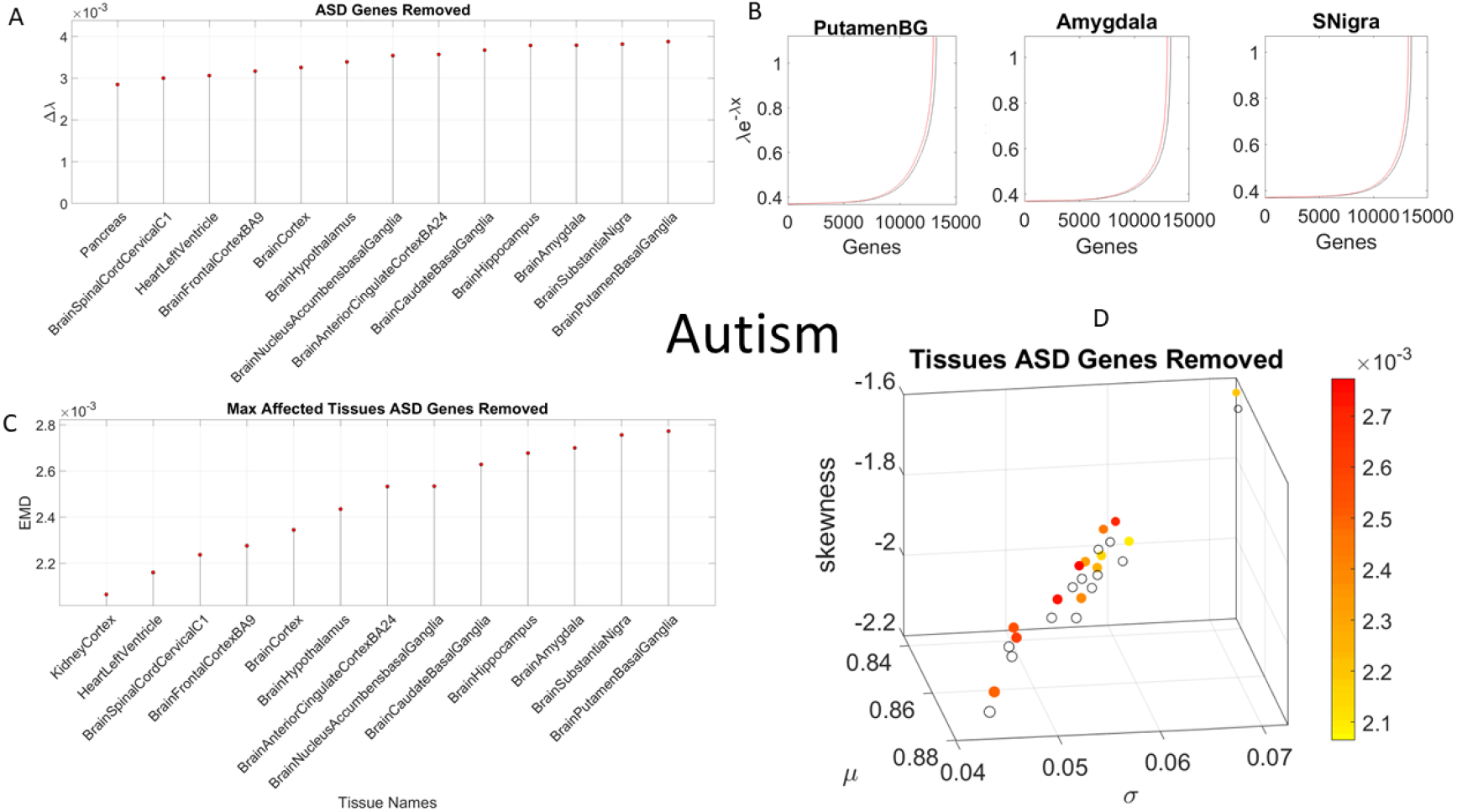
Removing the SFARI Autism genes from the normative human set reveals that the most affected tissues are those involved in neuromotor control, adaptation/learning, regulation and coordination. (Basal Ganglia, Substantia Nigra), emotion (Amygdala) and memory (Hippocampus.) (A) Highest median-ranked set of most affected tissues according to Δ*λ* quantifying the difference in exponential distributions describing the stochastic signatures of each tissue based on genes’ count (TCM). (B) Shift in the exponential description of the counts across genes for selected converging tissues common to Autism and all the other disorders under examination (other tissues shown in Table 1). (C) Consistent results are obtained using the continuous Gamma family of probability distributions empirically estimated using MLE for each tissue and EMD as a similarity metric to quantify departure from normative signatures upon SFARI Autism genes removal. (D) Visualization of the stochastic shift on the Gamma parameter space are depicted using a color gradient to represent the intensity of the shift and circles to represent the position (mean, variance, skewness and kurtosis proportional to the marker size), with open circles as the normative data and colored filled ones as those resulting from removing the SFARI Autism genes. Tissues are those revealed by the maximal genes’ expression changes in (A) and (B).

Figure 2 also depicts the results according to various stochastic metrics for the SFARI Autism genes removal. These include, the abovementioned MLE of the lambda rate parameter of the Exponential distribution characterizing the genes expression (count TPM) in Figure 2A. They also include the top median ranked group of tissues that are maximally affected according to the MLE of the continuous Gamma family of distributions that best characterizes the genes’ expression in the tissues. The shift in the Gamma parameters’ denoting the stochastic signatures for each top-ranked tissue is measured by the Earth Mover’s Distance (EMD) in Figure 2C, with the visualization of this Gamma parameters’ shift in Figure 2D. Figure 3 uses the same format as Figure 2 to show the outcome of removing the combined genes from the Ataxias, X-, FX-,PD and Mitochondrial disease (column 2 of Table 1) with genes reported in the literature (Table 2 of Methods section providing the numbers for genes and the sources.) The Supplementary Figures 1-3 show the results from selectively removing the genes for each of the neurological conditions of known genetic origins used in the combined case.

**Figure 3.**
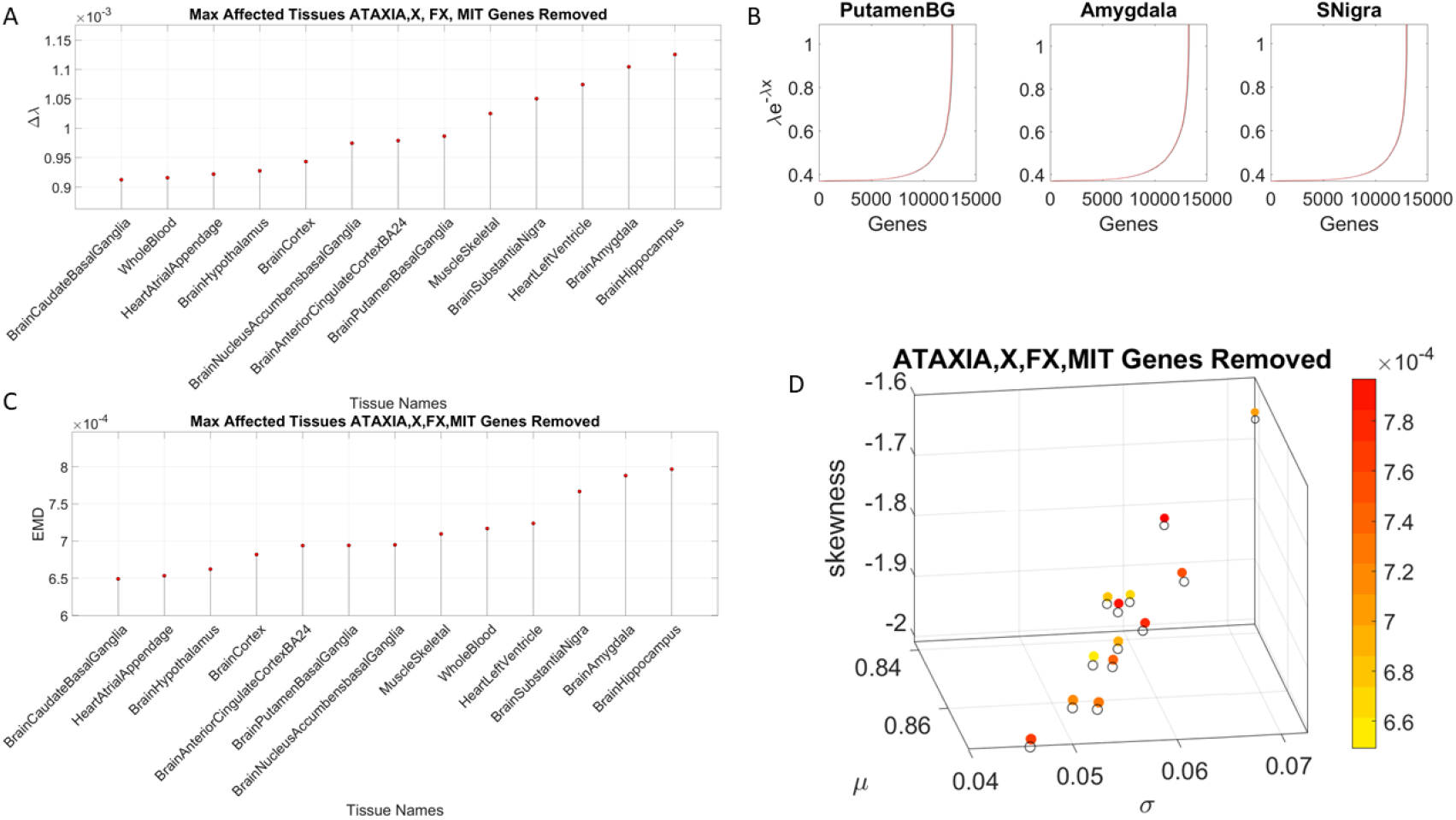
Removing the Ataxia, X, FX, PD and Mitochondria genes combined from the normative human set reveals that the most affected tissues are also involved in memory, neuromotor control, adaptation/learning, regulation coordination and autonomic heart function. (A)-(D) are as in Figure 2.

**Table 2.** Overlapping genes between the Ataxias (dominant and recessive genes) and the Parkinson’s disease with the genes from the SFARI Autism database. Scores in parenthesis refer to the scoring of the gene according to the SFARI Autism system (see Methods for explanation on each category). Syndromic is (4). Supplementary Figures 5-18 provide the GTEx violin plots of these genes’ expression in the top-ranked tissues unveiled by our analyses in Table 1. Supplementary Table 2 compiles the genes’ additional information from various sources in the clinical literature.

| SFARI Autism & Ataxia Dominant | SFARI Autism & Ataxia Recessive | SFARI Autism & (Early Onset) PD |
| --- | --- | --- |
| CACNA1C (1) | KCNJ10 (2) | RAB39B (3) |
| SCN2A (1) | WWOX (2) |  |
| ATP1A3 (3S) | GRID2 (3) |  |
| CCDC88C (3) | LAMA1 (3) |  |
| ITPR1 (3) | PEX7 (3) |  |
|  | SYNE1(3S) |  |
|  | CYP27A1 (4) |  |
|  | SNX14 (4) |  |

In all cases, the computation of the EMD differences revealed consistent patterns as those of the Δ*λ*-ranking. Moreover, the shifts of the signatures for each tissue in the Gamma parameter space clearly depict the departure from normative stochastic signatures for those tissues maximally ranked to be affected. These are shown in the panel D of Figures 2–3 using both a color code reflecting the EMD similarity metric and the stochastic Gamma parameter position, relative to the normative state where all genes are included. The summary of the genes’ expression on the 54 tissues from selectively removing the Autism, Ataxias, X, Fragile X, Parkinson’s disease and Mitochondria genes from the GETx set are shown in Figure 4. Supplementary Figures 1-3 show these results separately for each neurological condition, while Supplementary Figure 4 shows the top ranked Δ*λ* tissues for the SFARI Autism genes according to their scoring system. There we note that removal of the SFARI Autism syndromic genes reveal maximal differences in tissues of organs with smooth and cardiac muscles, linked to involuntary and autonomic function in the proposed taxonomy of Figure 1B.

**Figure 4.**
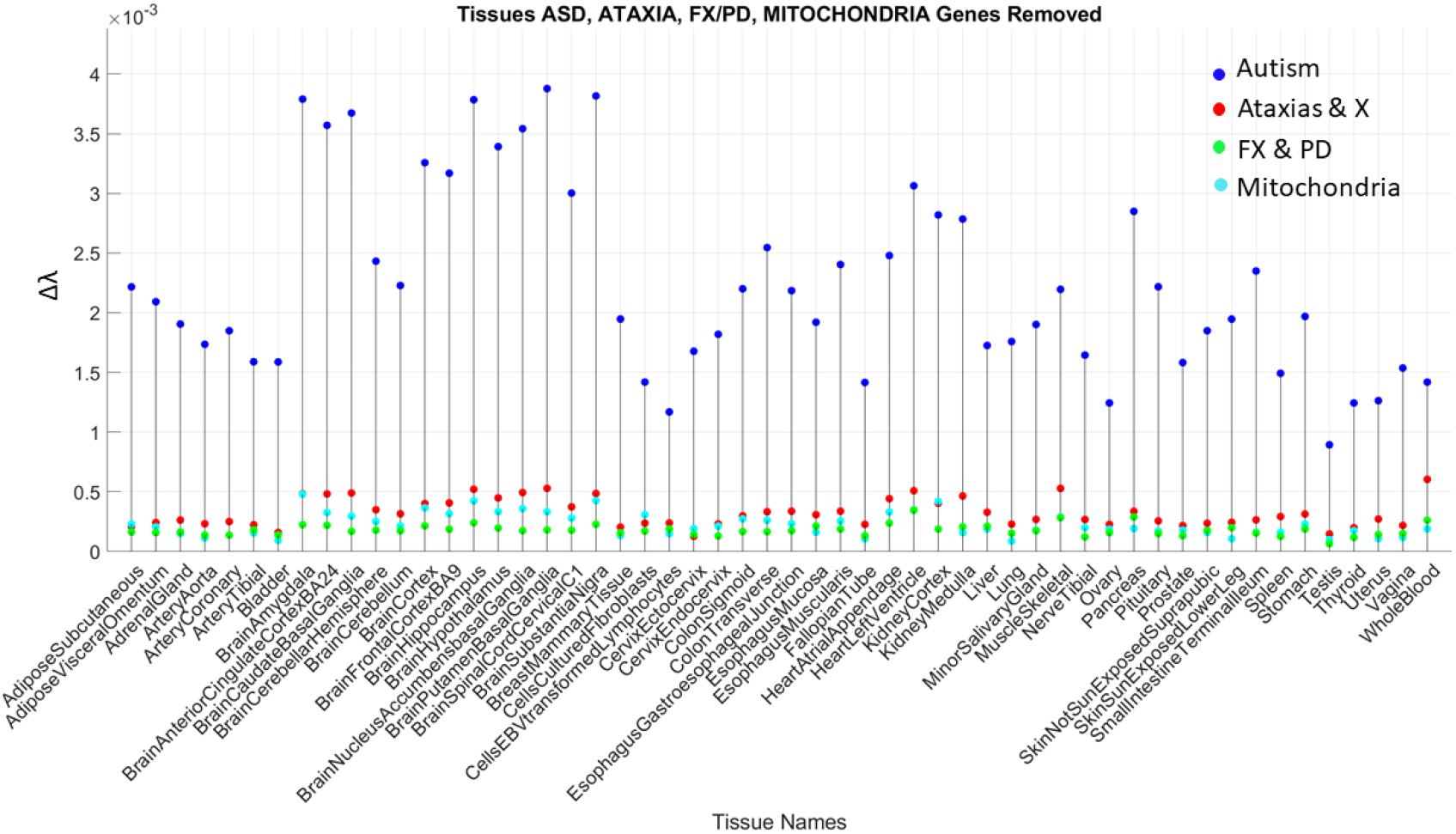
Selectively removing Autism, Ataxias, X, Fragile X, Parkinson’s disease and the Mitochondria genes from the normative human set in GTEx reveals that the most affected tissues are those involved in the nervous systems functioning, from voluntary, involuntary and autonomic levels of functional control, a result amenable to follow the proposed phylogenetically orderly taxonomy of the nervous systems maturation and function. The 54 tissues are displayed in alphabetical order.

Given the congruence between the tissues maximally affected by removing the SFARI Autism genes from the GTEx database, we next ascertain the extent to which these genes overlap with those used from the Ataxias in the literature. To that end, we divide them into the autosomal dominant, the autosomal recessive and the X-Chromosome genes. Figure 5A shows the result of this interrogation broken down by the Ataxia’s autosomal dominant, autosomal recessive and X-Chromosome genes. The genes listed in the figure overlap with those from the SFARI Autism genes. Table 2 lists them with the scoring from the SFARI Autism genes. The Supplementary Material Table 2 lists the phenotypic information of the disorders associated with these genes, as described by the clinical literature. Figure 5C lists the PD gene that overlaps with the SFARI Autism genes.

**Figure 5.**
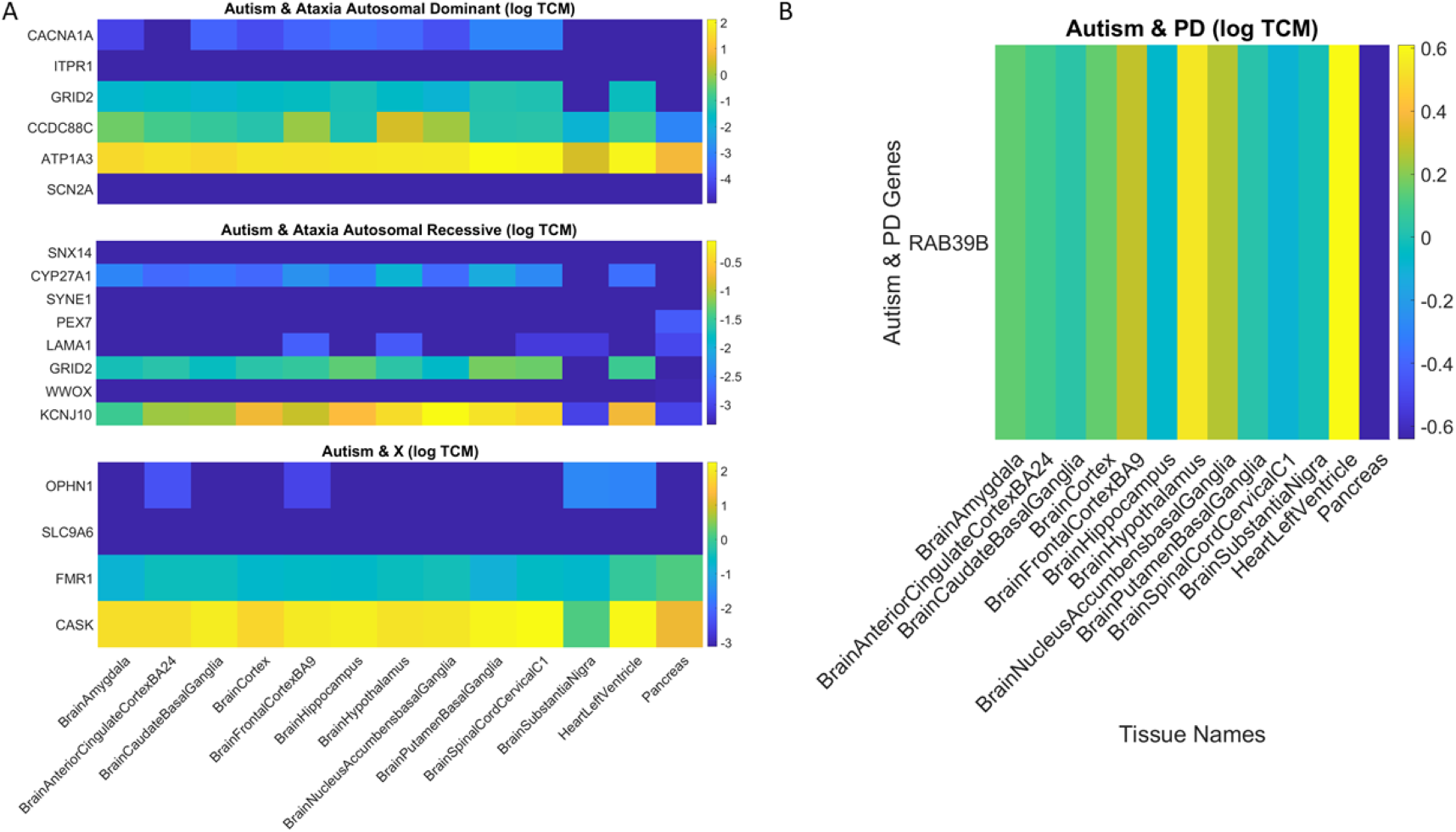
Overlapping genes between SFARI Autism set and the Ataxias (dominant and recessive and X-Chromosome) from the literature (A) and (B) from Parkinson’s disease. Color code is the gene expression (log TPM for each set.) Horizontal axis is the tissues’ names and vertical axis the genes’ names.

We note that removing this subset of 14 overlapping genes from the SFARI Autism set, does not change the primary result whereby the most affected tissues upon removal of the SFARI Autism set from the GTEx dataset are those involved in motor control, adaptation/learning, regulation, coordination and autonomic function, as well as memory and emotion. This is shown in Figure 6A-B and in the third column of Table 1. We also plot in Figure 6C the FX genes from the SFARI database for the top ranked tissues affected by the removal from the GTEx dataset of the SFARI Autism genes (i.e. without these FX genes.) Supplementary Figures 19-22 further provide details on X-Chromosome genes implicated in Autism according to the SFARI genes module.

**Figure 6.**
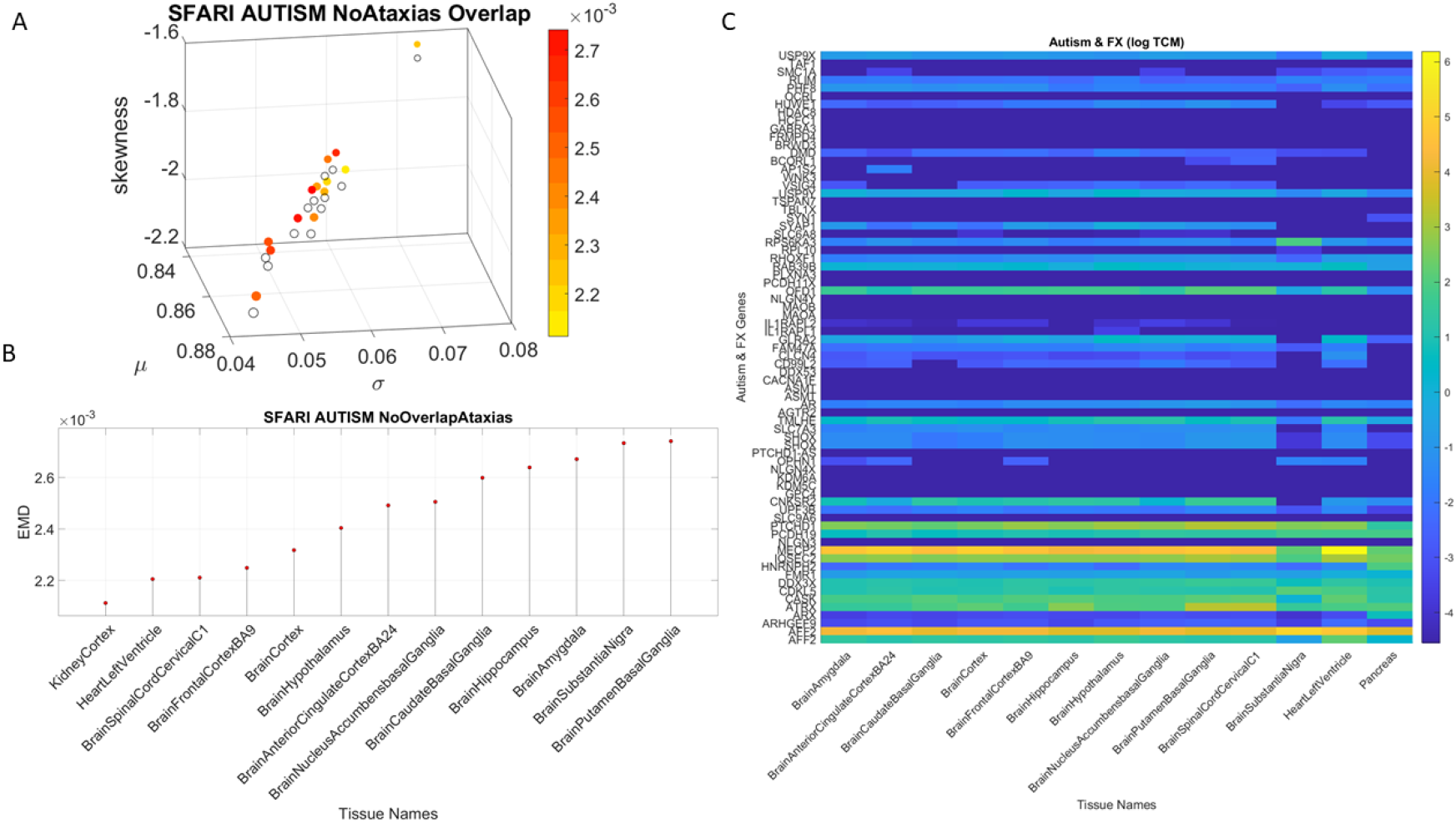
Effects on the tissues remain upon removal of the overlapping genes listed in Table 2 and Figure 5 from the SFARI Autism database. (Plots are as in Figures 2–3) (A) Gamma parameters’ shift and color coded by EMD denoting the difference between tissues with full genome *vs*. tissues upon removal of SFARI Autism genes minus those overlapping in Table 2. (B) Plots of the top-ranked tissues in (A) according to Δ*λ* top and EMD bottom. (C) FX genes set from SFARI genes expressed in the top-ranked affected tissues of Figure 5 and Table 1, *i.e.* those top-ranked tissues obtained upon removal of the SFARI Autism genes.

We find that common to all these well-known neurological conditions and to Autism is that the most affected tissues are the parts of the brain commonly linked to neurological disorders affecting the control and regulation of motions, emotions, and memory, followed by tissues affecting the autonomic level and lastly those affecting the involuntary smooth-muscle related tissues. As such, and based on growing evidence on disorders of voluntary, involuntary, and autonomic biorhythmic motions in autism, we propose to ***redefine*** the criterion to detect, treat and research autism as one that fundamentally describes disorders of the nervous systems. There are several advantages to this approach in view of new technological advances in Neuroscience, Genomics and Wearable Biosensing technologies, and the potential benefits that it could bring to affected individuals and their families.

This work used the SFARI genes module. This module implicitly relies on the clinical autism diagnosis criteria. These criteria have sidelined nervous systems issues in Autism in favor of *gold standards* that deem the autistic traits as socially inappropriate behaviors. Consequentially, autism neurology is virtually unknown officially. That is, no accreditation system in Psychology or Psychiatry includes these neurological criteria. Clinicians accredited to diagnose and treat autism do not have to acquire this knowledge base, nor do they have to update it with Continuing Education Units (CEUs.) Yet, we show here that there is high overlapping between well-known neurological disorders of known genetic origins and those in the SFARI Autism set. This result links Autistic behaviors with their underpinning neurobiology in very precise ways that may aid begin this new neurobiological roadmap for genetic inquiry in Autism.

We selectively removed genes from the human genome and assessed the extent to which removing those genes and treating the genes’ expression (counts TPM) in the tissues as a random process, would be conducive of phenotypic features aligned with those of well-known neurological disorders of genetic origins. Even upon removal from the genes of the SFARI set of the 14 Ataxia and Parkinson’s disease overlapping genes, and even though the genes’ expression on the tissues was treated as a random process (well-characterized by the memoryless Exponential distribution), at the basic tissue level, we found convergence between Autism and these disorders of the nervous systems.

To aid in connecting the layer of behaviors and the genome layer of the knowledge network, we used our proposed phylogenetic orderly taxonomy of nervous systems functions (Figure 1B), mapping nervous system’s functions to fundamental types of muscles defining voluntary, involuntary and autonomic levels of control. We simulated the possible worst-case scenario whereby for each of these disorders, the known genes were removed from the human full genome in the GTEx database. The results yielded, in all cases, tissues related to movement control, regulation and coordination, the very aspects of autism that clinical criteria selectively exclude (see Supplementary Material). They also reveal brain areas related to emotions and memory, and tissues linked to autonomic (cardiac) function as the most affected ones. Within the top ranked groups, we also found tissues present in organs with smooth muscle – vital for the overall function of the organism.

Unquestionably, somatic sensory motor issues are not only underscored by growing evidence in the field of Neuroscience [13, 15–17, 50]. They are also reported by self-advocates and parents. Yet, no framework existed that could link the genomic piece to that well established autism phenotype, so frequently reported in the neuroscientific literature [8, 11, 12, 41–45, 47–49, 51–54]. Up to now, the clinical criteria highlighting social deficits (as set by their expectations and assumptions) was the only link to the Autism genetics, via the behavioral-observational diagnosis (*e.g.* the ADOS test and the DSM criteria), but this gap between subjective observation and physical evidence had prevented the design of targeted treatments using a personalized approach [55].

Clinical criteria are much too coarse, so it inevitably gives rise to a broad, highly heterogeneous spectrum and offers no roadmap to automatically detect subtypes or to track dynamic age-dependent changes in the nervous systems. Using the proposed approach combining objective digital behavioral criteria and tissue-based genomic information tailored to each person’s genome, we may be able to automatically build subtypes of this condition and narrow down the criteria for treatments according to the affected functional levels of the nervous systems. Moreover, our analyses show that we will be able to do so independent of the clinical label.

At present, the “*one size fits all*” approach to autism diagnosis and treatments stresses the person nervous systems even more than it already is. The subjective behavioral criteria and the pipeline that it creates, starting with the diagnosis and following up with treatment recommendations to reshape assumed inappropriate behaviors, denies to the affected person the most basic right to autonomy and volitional control. Even the most rudimentary capacity for social readiness, which all autistics have, and the potential to flourish as an active member of our society, is abolished by the current behaviorist model of autism. But more worrisome yet, is the fact that by systematically ignoring the potential neurological and medical issues that disorders of the nervous systems give rise to, this current model denies the person the most basic human rights that life warrants.

The new model of Precision Medicine for Autism that we propose here offers the possibility to connect the multiple layers of the knowledge network and finally bridge behaviors to genomics. Furthermore, the present work linking Autism to the Ataxias, FX, Parkinson’s disease and Mitochondria disorders offers a fresh view of Autism that can help us learn from those other fields and advance the design of targeted treatments that are already ongoing in some of them. At the very least, we could follow the treatments. Such treatments would help mitigate medical issues present in a large sector of the autistic population since birth. And because of their personalized nature, they would (1) not be arbitrarily imposed on those who do not need them and (2) be adaptable to the age-dependent shifting needs.

When we started this line of inquiry, we already knew that Autism had movement sensing differences impacting social exchange as we know it [56]. We also knew of the competence of Autistic individuals, of their learning predispositions, preferences, and capabilities. What we did not ever suspect was that a random process such as the gross removal of the SFARI Autism genes from the full human genome was going to result in such convergence with the well-known neurological disorders of genetics origins. This work offers a new roadmap and a unifying enabling approach for autism research, diagnostics, and personalized targeted treatments design.

## Materials and Methods

We combine the data sets from the genes scoring module of the Simons Foundation Autism Research Initiative (SFARI) and from the GTEx Portal human RNA-Seq (Transcripts Per Million TPM)^1^ specifically using the files denoted in the Appendix B. In addition to the human genes and the autism genes (scored by SFARI), the ATAXIA genes, the X-genes and the FX-genes were taken from various literature reviews tabulating the hereditary Ataxias [57, 58] and the mitochondrial disorders [59]. Genes identified in Parkinson’s disease were taken from [60–65].

The SFARI Autism categories that we used were those as of 03-04-2020. Quoting from their site:

- CATEGORY 1 Genes in this category are all found on the SPARK gene list. Each of these genes has been clearly implicated in ASD—typically by the presence of at least three de novo likely-gene-disrupting mutations being reported in the literature—and such mutations identified in the sequencing of the SPARK cohort are typically returned to the participants. Some of these genes meet the most rigorous threshold of genome-wide significance; all at least meet a threshold false discovery rate of < 0.1.
- CATEGORY 2 Genes with two reported de novo likely-gene-disrupting mutations. A gene uniquely implicated by a genome-wide association study, either reaching genome-wide significance or, if not, consistently replicated and accompanied by evidence that the risk variant has a functional effect.
- CATEGORY 3 Genes with a single reported de novo likely-gene-disrupting mutation. Evidence from a significant but unreplicated association study, or a series of rare inherited mutations for which there is not a rigorous statistical comparison with controls.
- SYNDROMIC The syndromic category includes mutations that are associated with a substantial degree of increased risk and consistently linked to additional characteristics not required for an ASD diagnosis. If there is independent evidence implicating a gene in idiopathic ASD, it will be listed as “#S” (e.g., 2S, 3S). If there is no such independent evidence, the gene will be listed simply as “S”.

The GTEx dataset is as of 06-05-2017 v8 release. For every gene in the Autism, Ataxia, X, FX, Mitochondrial diseases, and Parkinson’s disease, we first confirmed the presence of the gene in the GTEx data set and then incorporated it to the analyses.

### Count Normalization

The GTEx matrix of RNA-seq genes (*56,146*) along the rows x the tissues (*54*) along the columns was transposed (*54* × *56,246*), such that we expressed each tissue as a function of the gene expression denoted by the count (TPM). Each individual count value was then normalized using equation (1)

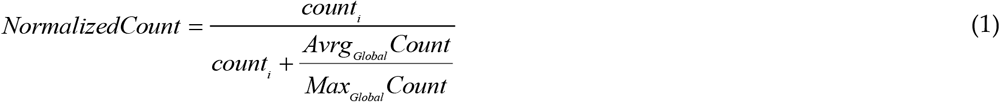

Here *count_i_* is the count value of the *gene_i_*, *Avrg_Global_Count* is the overall average of the matrix of values taken along the columns and the rows. *Max_Global_Count* is the maximum count value, also taken globally across the matrix values. Figure 8 shows the original count numbers (Figure 7A) and the normalized version (coined Micro-Movement Spikes MMS) in Figure 7B. Figure 7C shows the MMS derived from the fluctuations in counts normalized by equation (1) while Figure 7D shows the histograms of the peaks (marked in red dots) for different tissues and genes scored by SFARI.

**Figure 7.**
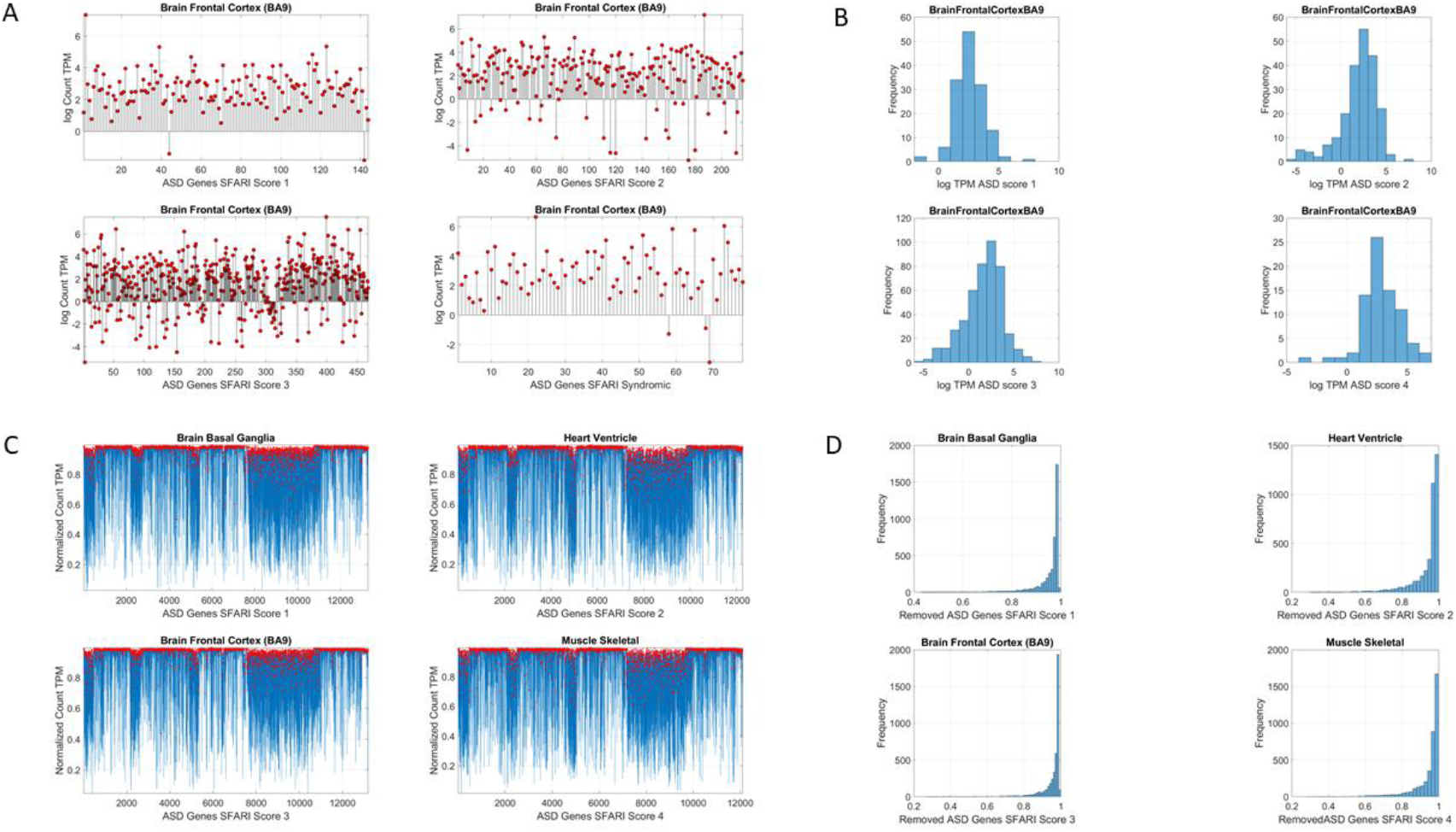
Analytical methods. (A) Sample raw data consisting of log count (TPM) for different scored genes expressed in the brain frontal cortex (Brodman Area 9.) (B) Histograms of the log Count TPM for each case in (A). (C) Upon removal of the SFARI Autism genes, micro-fluctuations’ spikes in the normalized count, with deviations taken relative to empirically estimated mean, global averaged count and global maximal count in Equation (1). (D) Histograms of the normalized micro-fluctuations’ spikes.

### Genes Removal

For each of the disorders of interest, Autism, Ataxias, X-chromosome, Fragile-X, Mitochondria, Parkinson’s disease, we remove from the human GTEx data set the genes from the SFARI Autism genes module, and/or from the literature relevant to each condition. We then treat the resulting count series as a random process. We use the exponential distribution to characterize it and to assess the differential expression across the tissues relative to no removal in the original human genome.

The question that we ask is, given the known neurological phenotypes of the known disorders (Parkinson’s disease, the Ataxias, the X-Chromosome, Fragile-X and Mitochondria), is there convergence between the most affected tissues (as measured by stochastic change as explained in the next section) of these known disorders (and their known neurological functions) and the changes in the tissues that will be obtained by removing the genes associated with Autism? Table 2 shows the number of genes removed in each condition.

**Table 2.** Genes distributions used in the removal process and literature sources.

| Condition | Number of Genes | Source |
| --- | --- | --- |
| Parkinson | 17 | Lit Review [60, 62-65] |
| Ataxia Autosomal Recessive | 70 | Lit Review [57, 58] |
| Ataxia Autosomal Dominant | 46 | Lit Review [57, 58] |
| X-Chromosome | 6 | Lit Review [57, 58] |
| Fragile X | 73 <sup>1</sup> | SFARI Genes Module |
| (SFARI FX scores) | (17,11,30,15) |  |
| Mitochondria | 41 | Lit Review [59] |
| Autism | 906 <sup>1</sup> | SFARI Genes Module |
| (SFARI Autism scores) | (144,216,468,78) |  |

<sup>1</sup> SFARI scores for Autism and FX genes were also used to assess each scored module separately

### Stochastic Analyses

Since the count values for each tissue can be conceived as a random series of numbers, we use maximum likelihood estimation (MLE) to model the numbers representing the counts, as generated by the exponential distribution using equation (2)

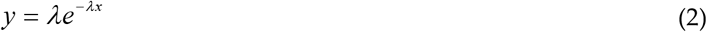

Here *x* represents the normalized count value (as per Equation (1)) and *y* the value from the exponential distribution. We seek the value of *λ* to model this counting random process representing the genes’ expression in the tissue. To that end, we estimate the likelihood *L*(λ|*x*_1_,*x*_2_,…,*x_n_*) where the series of counts *x_i_*, with *i* ranging from *1* to *n* represent the normalized counts (according to Equation 1) across all genes for one tissue. Appendix B shows the steps to find *λ*. And this is computed for each of the 54 tissues. We then rank the departure of *λ* (resulting from genes removal) from the *λ* obtained for the full human genome (see below).

### Stochastic Analyses – Visualization of change relative to the normative data of the full human genome

Using MLE we also obtain for each of the 54 tissues, the frequency histograms of the normalized counts across all genes and fit the continuous Gamma family of probability distributions with shape (*a*) and scale (*b*) values, to obtain the Gamma moments and plot them on a parameter space. We do this to visualize the spread of the tissues and their shift upon genes removal. To that end, we plot the mean, the variance, and the skewness across the x-, y- and z-axis respectively. We plot the size of the marker representing the tissue proportional to the kurtosis value, and we color the marker based on the change relative to the original genome count (*i.e.* containing all the genes, without removal.)

To measure the stochastic shift between the tissues from the full genome and those upon removal of the genes identified with each known neurological condition, we use the Kantarovich-Wasserstein distance [66–68], also known as the Earth Mover’s Distance [69]. This distance metric quantifies the amount of work that it takes to transform one frequency histogram into another. In this case, the histograms in question are *e.g.* the histogram of the genes counts for a given tissue obtained from the full genome *vs.* that of the counts upon removing the genes associated with the disorder (*e.g.* ASD).

Once we obtain the differences using the EMD metric, we rank the tissues, sorting from minimal to maximal departure (Δλ) of the modified genome from the original set, using the *λ*-values obtained from the tissues in the neurotypical genome containing the counts from all the genes. We then compare the Δ*λ* across these disorders, detecting which tissues are maximally affected by the gene’s removal. To that end, we compute the median Δ*λ* and rank all tissues above and below the median value. We then, for each block median-rank again and obtain 4 blocks thus ranked. We examine the top ranked block and identify the tissues revealed as the maximally affected ones by the removal of the genes from the GTEx full set. We also do this using the EMD derived from the Gamma fit for visualization. The highest ranked group is then compared across all conditions, ASD *vs.* those neurologically defined. We annotate the neurological functions that such tissues are known to maximally disrupt. And we ask if in the case of ASD, there is convergence between the tissue outcome, upon removing the SFARI genes from the human genome-tissue model, and the outcome upon removing those genes tied to the other known disorders of the nervous systems.

## Patents

EBT holds the US Patent “Methods and Systems for the Diagnoses and Treatments of Nervous Systems Disorders” combined in the paper as micro-movement spikes, MMS data type and Gamma process.

## Author Contributions

Conceptualization, methodology, software, validation, formal analysis, investigation, resources, data curation, writing EBT.

## Acknowledgments

I thank the SFARI researchers for the curation and maintenance of their genes module and the compilation of literature database supporting the repository.

## APPENDIX A

We estimate the likelihood *L*(λ|*x*_1_,*x*_2_,…,*x_n_*) where *x_i_* is the series of counts representing the gene expression on each given tissue and *i* ranges from *1* to *n*, the number of genes.

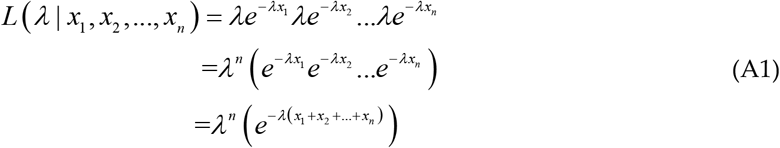

To obtain the maximum likelihood, we take the derivative of the likelihood in equation (A1) and set it to 0 (since the derivative is 0 at the maximum likelihood value).

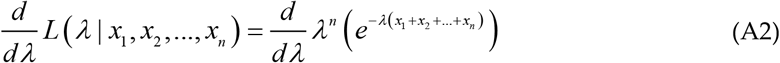

We take the *log* here because the derivative of the function and the derivative of the *log* of the function equals *0* at the same point, so for the purposes of finding where the derivative is *0*, the original function in equation (A2) and the *log* of it are interchangeable.

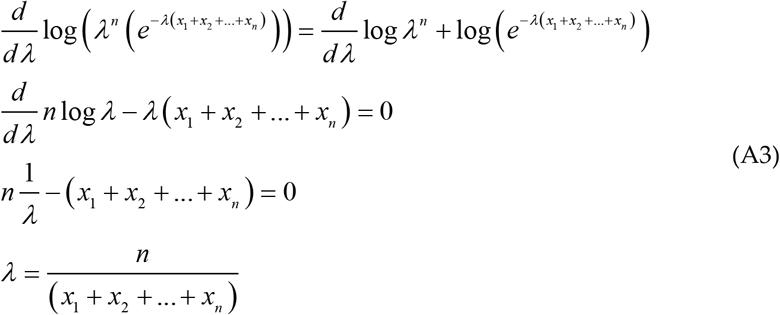

And with this result in equation (A3), we can obtain the maximum likelihood estimate of each *λ*_*j*_, given all the 56,146 genes expressed with some random value for each of the *j* = 1: 54 tissues.

## Appendix B

The data file name from the GTEx Portal https://www.gtexportal.org/home/datasets used in this paper is GTEx_Analysis_2017-06-05_v8_RNASeQCv1.1.9_gene_median_tpm.gct.csv

The data file name from SFARI Gene is located at https://gene.sfari.org/database/human-gene/ and named SFARI-Gene_genes_03-04-2020release_03-05-2020export.csv

© 2020 by the authors. Submitted for possible open access publication under the terms and conditions of the Creative Commons Attribution (CC BY) license (http://creativecommons.org/licenses/by/4.0/).

## Footnotes

1 Taken from the site: “Transcripts Per Million (TPM) is a normalization method for RNA-seq, should be read as *for every 1,000,000 RNA molecules in the RNA-seq sample, x came from this gene/transcript*. For each transcript in the gene model, the number (raw count) of reads mapped is divided by the transcript’s length, giving a normalized transcript-level expression. The distribution of ambiguous reads (between transcripts of the same gene, or between different genes) is handled by OmicSoft’s RSEM implementation. The sum of ALL normalized transcript expression values is divided by 1,000,000, to create a scaling factor. Each transcript’s normalized expression is divided by the scaling factor, which results in the TPM value. Gene-level TPM’s are calculated by summing up the transcript-level TPM for each gene. In this scaling, the sum of all TPMs (transcript-level or gene-level) should always equal 1,000,000. For cells that have approximately the same number of transcripts-per-cell, the TPM expression values can be compared between these cells to estimate relative abundance. For a given sample, TPM values will linearly scale with FPKM values for genes or transcripts, but FPKM will not add up to 1,000,000, so TPM can also be thought as FPKM, scaled to sum to 1,000,000.”

